# Vaccinomics Approach for Scheming Potential Epitope-Based Peptide Vaccine by Targeting L-protein of Marburg Virus

**DOI:** 10.1101/2020.12.29.424691

**Authors:** Tahmina Pervin, Arafat Rahman Oany

## Abstract

Marburg virus is one of the world’s most threatening diseases, causing extreme hemorrhagic fever, with a death rate of up to ninety percent. The Food and Drug Administration (FDA) currently not authorized any treatments or vaccinations for the hindrance and post-exposure of the Marburg virus. In the Present study, the vaccinomics methodology was adopted to design a potential novel peptide vaccine against the Marburg virus, targeting RNA-directed RNA polymerase (L). A total of 48 L-proteins from diverse variants of the Marburg virus were collected from the NCBI GenBank server and used to classify the best antigenic protein leading to predict equally T and B-cell epitopes. Initially, the top 26 epitopes were evaluated for the attraction with major histocompatibility complex (MHC) class I and II alleles. Finally, four prospective central epitopes NLSDLTFLI, FRYEFTRHF, YRLRNSTAL, and YRVRNVQTL were carefully chosen. Among these, FRYEFTRHF and YRVRNVQTL peptides showed 100% conservancy. Though YRLRNSTAL showed 95.74% conservancy, it demonstrated the highest combined score as T cell epitope (2.5461) and population coverage of 94.42% among the whole world population. The epitope was found non-allergenic, and docking interactions with human leukocyte antigens (HLAs) also verified. Finally, in vivo analysis of the recommended peptides might contribute to the advancement of an efficient and exclusively prevalent vaccine that would be an active route to impede the virus spreading.

## 1.0 Introduction

A member of the genera Marburg virus (MABV) belongs to Filoviridae family that causes an acute life-threatening hemorrhagic fever in human being and other primates. The fever caused by MABV is identical to the Ebola virus (EBOV) disease and therefore is characterized by fever, severe inflammatory reaction, pathological coagulation, and systemic hemorrhaging [1]. To date, Marburg virus disease (MVD) is related to 469 total cases and 376 reported deaths [2, 3]. Though fewer cases for MABV recorded, forthcoming epidemics and the rapid expansion of MABV to non-endemic areas are of boundless distress. The mortality rate of this virus reported about 81 percent[1].

The Egyptian rousette *(Rousettus aegyptiacus)* is publicly classified as the Marburg virus-host and has a wide geographical distribution. The pretended cause for the dispersal of MVD might be the close contact between humans and animals like, other than human primates, bats, and cattle [4]. It has been listed by World Health Organization (WHO) as a category 4 risk agent because of its lethality [5].

The Marburg virus possesses about 19 kilobase-pair long, negative sense, un-segmented RNA genome, encoding 7 open reading frames (ORF); nucleoprotein NP, virion protein (VP) 35, VP40, glycoprotein (GP), VP24, and viral RNA-dependent RNA polymerase (L) [6]. The whole-genome of filovirus is enveloped into a single threadlike virion, having 790 to 970 nm length and 80 nm width [7]. Based on the structure and functions of Marburg virus L-protein, the enzymatically active subunit (L) of the MARV polymerase comprises of 2331 amino acids [8]. The L-protein conjunction with VP 35, the polymerase cofactor, forms the RNA-dependent polymerase complex that is crucial for transcription and duplication of the virus [9].

Because of the emergence of the Marburg virus outbreak, novel therapeutic targets against this pathogen need to be identified immediately. Currently, no FDA approved vaccines or drugs are available to defend the human against the Marburg virus [10]. The only prime treatment of patients during the epidemics was sympathetic care (fluids, antimicrobials, blood transfusion) [11, 12].

Identifying precise epitopes from pathogens causing infections has ominously advanced the progress of peptide vaccines grounded on the epitope. Better knowledge on the bio-molecular base of target identification and human leukocyte antigen (HLA) binding motives, culminated in the development of logically engineered vaccines based uniquely on algorithms forecasting the binding of the epitope to HLA [13–16]. The epitope-based vaccine is chemically constant, more precise, and without any potentially pathogenic or oncogenic exposure [17, 18], but the formulation of laboratory-based aspirant epitope is not only costly but also painstaking, requiring diverse laboratory medicine experiments for ultimate epitope choice.

Vaccinomics means the implementation of combined expertise from various fields, including immunogenetics and immunogenomics, to establish and understand the immune response of candidates for the next generation vaccine [19]. At present, different vaccinomics databases are available for identifying particular B lymphocyte epitopes and highly sensitive and specific HLA ligands [20, 21]. The vaccinomics strategy has already proved its promise with optimal findings in the detection of the retained epitope for human coronavirus [14], Ebola virus [16], and *Shigella* [15].

In the present work, vaccinomics approaches have been executed to design potential conserved epitope candidate that can be used for the vaccine origination against the deadly Marburg virus, by targeting protein L, with an anticipation of further wet lab endorsement.

## 2.0 Methods

The overall schematic diagram for vaccine designing is shown in **Figure 1**.

**Figure 01:**
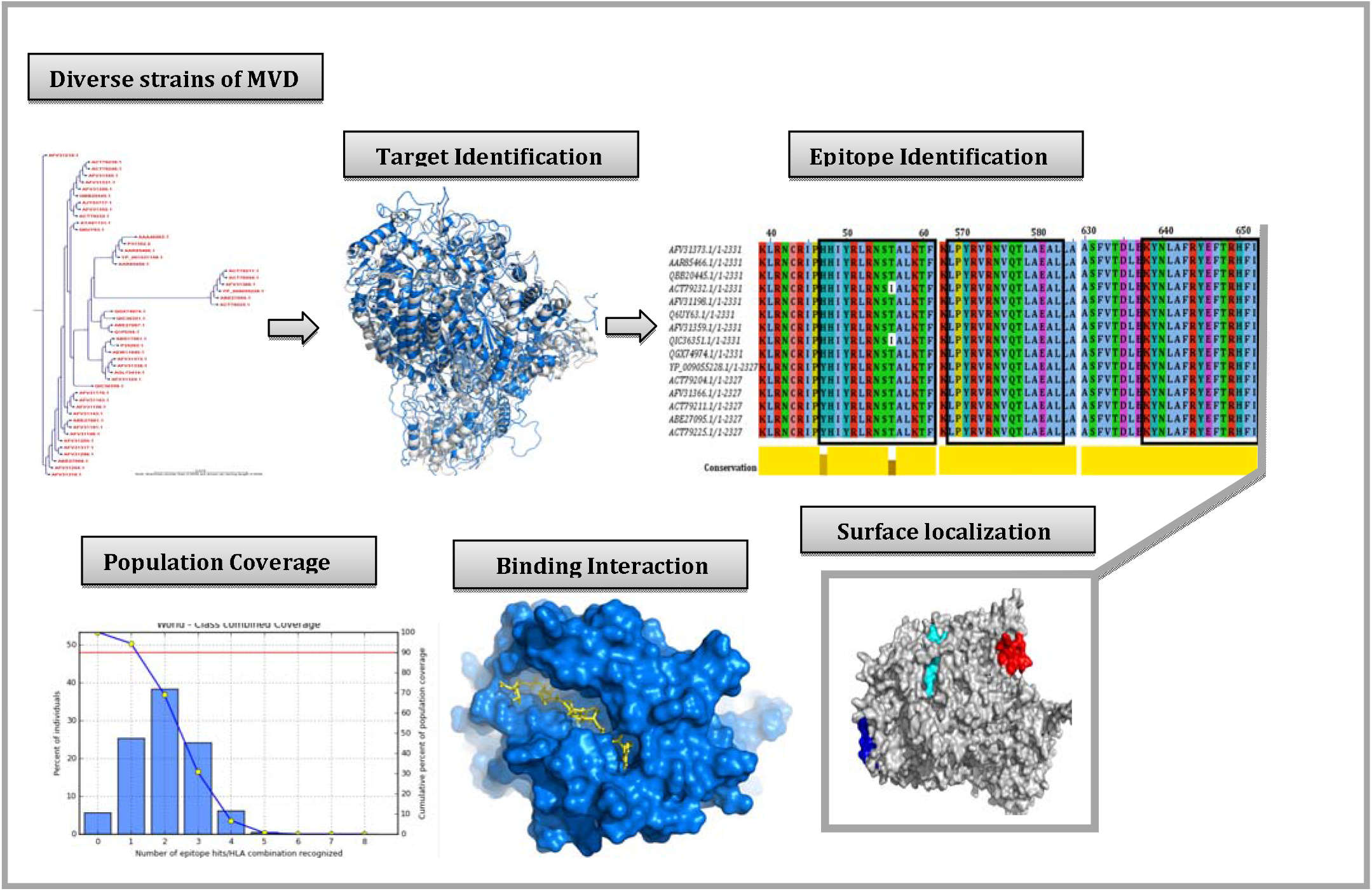
The overall schematic diagram for vaccine designing.

### 2.1 Sequence collection and antigenic protein determination

The L-protein sequences of the Marburg virus were collected from GenBank [22] and evaluated by the Immunomedicine Group server based on the Kolaskar method [23], to ascertain the maximum effective antigenic protein.

### 2.2 T-Cell Epitope Extrapolation and Affinity with MHC

The epitope estimation for the corresponding protein and its affinity score with MHC class I and class II allele were determined using the previously used approach [14, 16]. Briefly, the NetCTL server [24] assessed the effective cytotoxic T-lymphocyte epitopes from the most immunogenic protein. The default algorithms were used for this assessment. Epitopes having maximum scores were selected.

The Immune Epitope Database (IEDB) T-cell epitope detection methods were utilized to determine an association with both MHC molecules [25] [26, 27]. To measure the half-maximum inhibitory concentration (IC_50_) for the binding of a pre-selected 9.0-mer epitope with MHC-I, the stabilized matrix method (SMM) was employed. The IEDB-recommended approach was used to study MHC class II interaction for the particular HLA-DP, HLA-DQ, and HLA-DR loci. Taking into consideration the pre-defined 9-mer epitope and its preserved region in the Marburg virus, fifteen-mer epitopes were intended for binding interaction analysis of MHC-II. In case of MHC-I, the epitopes consisting of IC_50_ < 250nM and the epitopes containing percentile rank <50 for MHC-II alleles were chosen for even more assessments.

### 2.3 Epitope Management and Population Distribution Study

The aspirant epitope preservation was checked using a web-based epitope conservation method available in the IEDB research database [28]. The degree of conservation of each possible epitope was evaluated by taking attributes from the database of all L-protein sequences of different strains. Additionally multiple sequence alignment (MSA) was constructed by BioEdit software [29] and retrieved through Jalview (http://www.jalview.org/) tool to identify the epitope locations inside the sequences. The phylogram was generated through the CLC Sequence Viewer [30]. Furthermore, the WebLogo server [31] was used to visualize the conserved region of the MSA. The population distribution for the epitope was analyzed by IEDB estimation tool [32], and the collective value for both MHC groups was calculated.

### 2.4 Homology modeling, energy minimization and structural authentication

The homology model of the targeted protein was acquired by MODELLERv9 [33] and the PROCHECK server of the SWISS-MODEL Workspace evaluated the projected structure [34] [35]. The best suitable template was used for the construction of the model to cover the whole region of the protein. Before the assessment, the energy minimization of the structure was done by the GROMOS 96 [36]force field. The superposition of the prototype and model configuration had been examined to ensure the model’s fair consistency. Additionally, the ProSA-web server [37, 38] was also utilized for assessing the model quality through Z-score measurement.

### 2.5 Molecular docking and association of HLA allele’s scrutiny

Studies of docking were also carried out using the best available epitope, adopting the method used in previous studies [13, 14]. Auto-Dock Vina [39] was used for the docking analysis. In our analysis we designated HLA-C*07:02 and DRB1*04:01 as the aspirants for MHC-I and II, respectively, for docking study since they are accessible hits in the Protein Data Bank (PDB) server. The structure 5VGE, HLA-C*07:02 complexes with RYR peptide, and 5JLZ- human HLA-DRB 1*04:01 complexes with modified alpha-enolase peptide- were extracted from the Research Collaboratory for Structural Bioinformatics (RCSB) protein databank [40]. Finally, the complex-structures were generalized through PyMOL (Version 1.5.0.4) for the ultimate docking analysis.

For epitope 3D structure transformation, the preferred PEP-FOLD server was utilized to convert the structure from the sequences [41]. To evaluate the relationship with HLA alleles, the 9-mer epitope of the MHC I and 15-mer epitope of the MHC II molecule were used. Finally, all the proteins and the peptides were minimized using GROMOS 96 [36] force field for the docking analysis through using Swiss-Pdb viewer [42].

The grid-line for the docking of MHC-I molecules was of X: 58.9062, Y: 28.6331, and Z: 30.7545 (Angstrom) at the center of X: 16.9834, Y: −61.6639, and Z: 17.8431 and the grid-line for the docking of MHC-II molecules was of X: 26.9773, Y: 33.5139, and Z: 26.0699 (Angstrom) at the center of X: −41.0931, Y: 9.7731, and Z: −26.5406. A control docking was also performed with an experimentally acknowledged peptide-MHC- compound. The human HLA-C*07:02 bound with RYR peptide (PDB ID: 5VGE) was selected for this study. The gridline for the docking focusing at X: 25.6546, Y: −60.4063, and Z: 11.8845.

### 2.6 Allergenicity Assessment and B-Cell Epitope Detection

A Web-based AlgPred server was used to estimate the allergenicity of the possible epitopes [43] by using the support vector machine (SVM) algorithm at a threshold value of −0.4. Estimating technique follows guidelines of the Food and Agriculture Organization/World Health Organization, 2003. IEDB-AR screened the projected T-cell epitope (15-mer) using multiple web-based methods for the feasibility of the B-cell epitope [23, 44–48].

## 3.0 Results

### 3.1 Sequence collection and antigenic protein determination

From the NCBI GenBank database, entirely 48 L- protein molecules from diverse varieties of the Marburg virus were obtained. The L-proteins MSA was retrieved from the BioEdit software via ClustalW using 1000 bootstrap replicates (**Figure S1**). The CLC Sequence Viewer had been utilized to construct the phylogram based on MSA obtained from BioEdit, to determine the distinction between the sequences received and is depicted in the **Figure 2**. Finally, after subsequent evaluation from Immunomedicine Group server, the maximum antigenicity score of 1.0392 was found for the accession number AFV31373.1 (**Table S1**), and subsequently analyzed for the highly immunogenic epitope.

**Figure 2:**
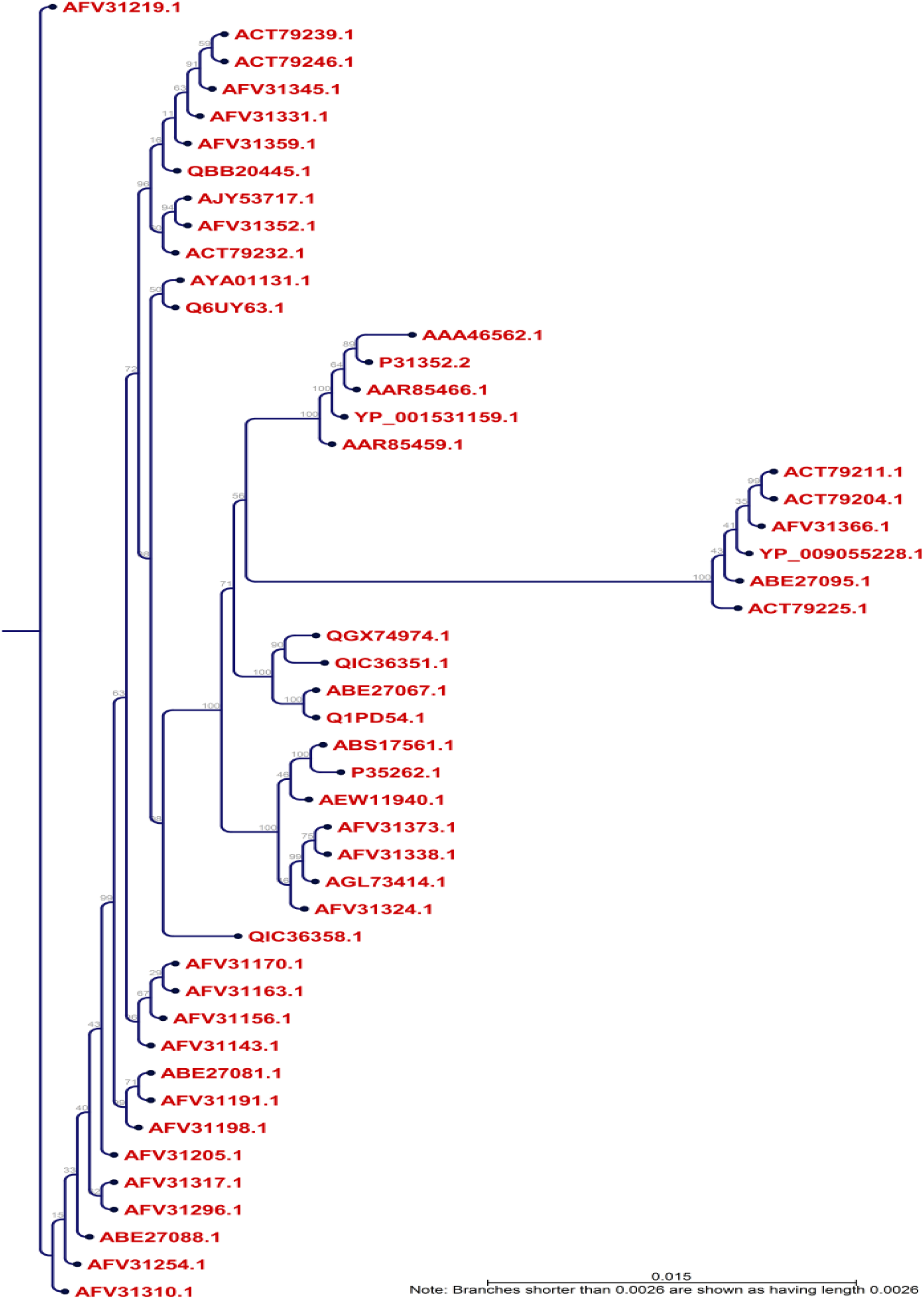
Phylogenetic tree showing the evolutionary divergence among the different RNA-dependent RNA polymerase-L proteins of the Marburg virus. Notes: Here, the phylogramic view is shown with appropriate distance among the different strains. The blue dotted view indicates the node of the tree.

### 3.2 T-Cell Epitope Extrapolation and Affinity with MHC

The NetCTLv1.2 server was utilized for the identification of T-cell epitopes where the calculation was restricted to 12 MHC-I supertypes. The top 26 epitopes (**Table 1**) were selected from the supertypes for further comprehensive analysis based on the cumulative values. **Table 2** lists the restricted MHC-I alleles for which the epitope exhibited increased specificity (IC_50_<250 nM) and in **Table 3** displays findings from the MHC-II interaction study (percentile rank < 50).

### 3.3 Epitope Management and Population Distribution Study

The IEDB conservation analysis method analyzed the epitope conservation of the anticipated epitopes as shown in **Table 4**. The positions of the predicted epitopes are shown in the MSA of L proteins (**Figure 3)**. Here, we used only 15 diverse sequences from the total retrieved proteins, 48 sequences, for the proper annotation. The entire world population coverage was calculated based on the combined MHC-I and MHC-II class with the top chosen interrelated alleles (**Figure 4** and **Table S2**).

**Figure 3:**
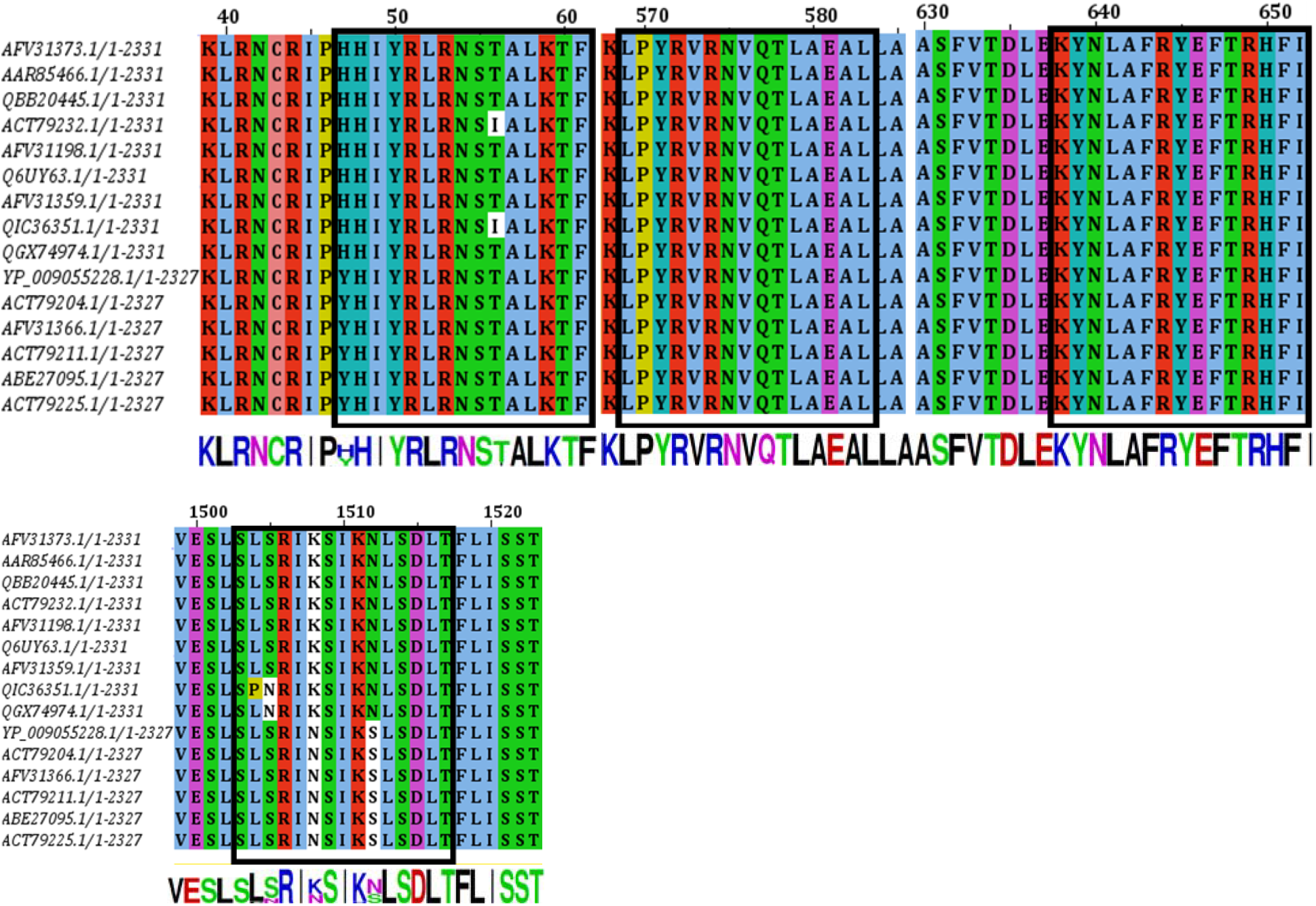
MSA-based location identification of the different epitopes within the L proteins of Marburg virus. Here the epitopes are shown by boxed (black) region within the MSA. The conservancies of the sequences are displayed by sequence to logo at the bottom of the total MSA. Here, only the best four epitopes, 15 mer including 9 mer, are shown.

**Figure 4:**
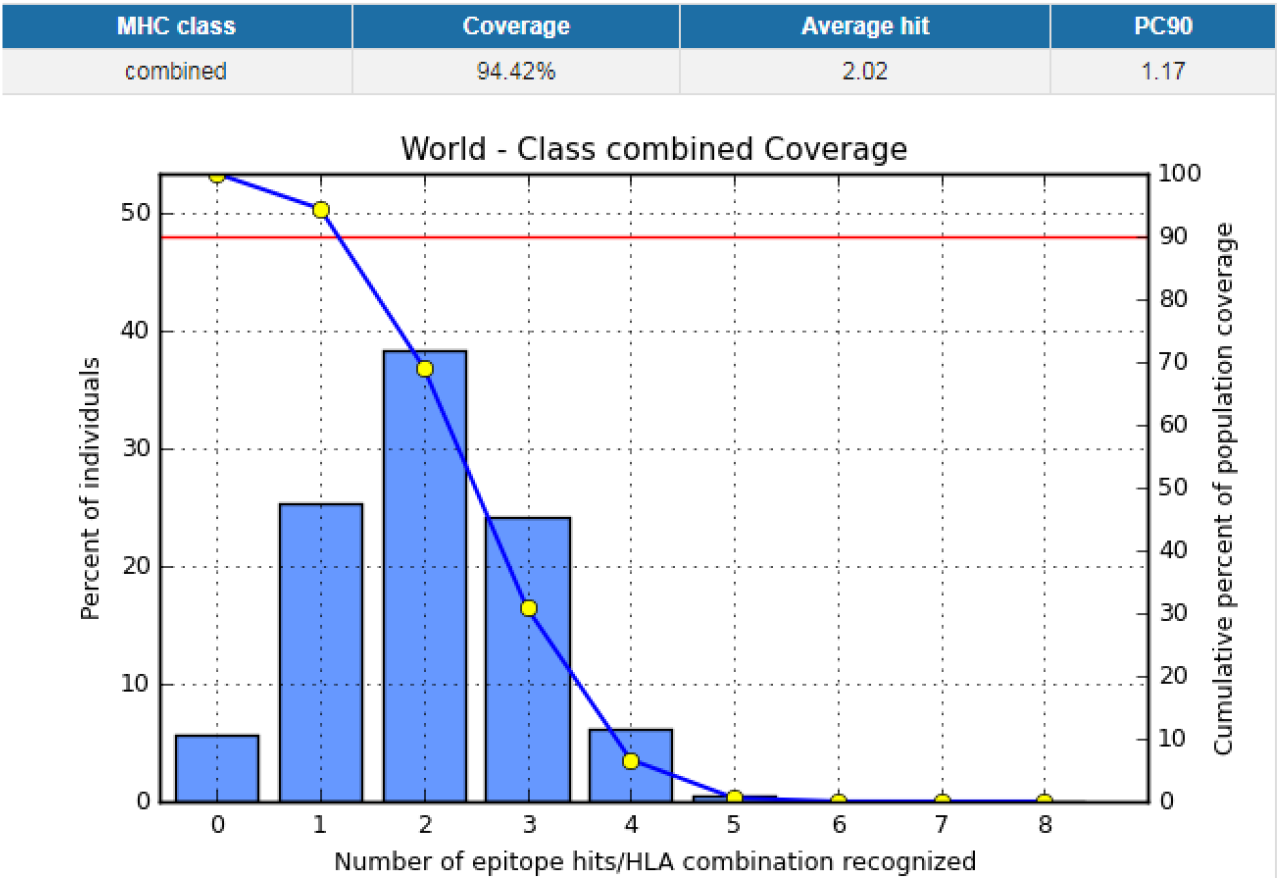
Population coverage analysis for the top predicted epitope (YRLRNSTAL) based on the HLA interaction. Here, the whole world populations are assessed for the proposed epitope. The combined prediction for both of the MHC has been shown. Here, the number 1 bar for all the analyses represents out-predicted epitope. Notes: in the graphs, the line (-o-) represents the cumulative percentage of population coverage of the epitopes; the bars represent the population coverage for each epitope.

### 3.4 Homology modeling, energy minimization and structural authentication

The three-dimensional configuration of the selected protein was constructed by MODELLER using the paramount template based modeling method and the template 6V85_A was utilized for this purposes. The energy minimization of the model was done from energy level 43319.410 to – 60879.637 through GROMOS 96 force field. The superposition view between the model and template is shown in **Figure 5a** with an RMSD of 2.08. The validation of the model was assessed by the PROCHECK server through the Ramachandran plot and is shown in **Figure 5c**, where 90.51% of amino acid residues were originated within the most-favored area. Furthermore, the ProSA Z score was also implemented for validation and shown in **Figure 5b**. Additionally, the top four epitopes were shown on the protein surface in **Figure 5c**.

**Figure 5:**
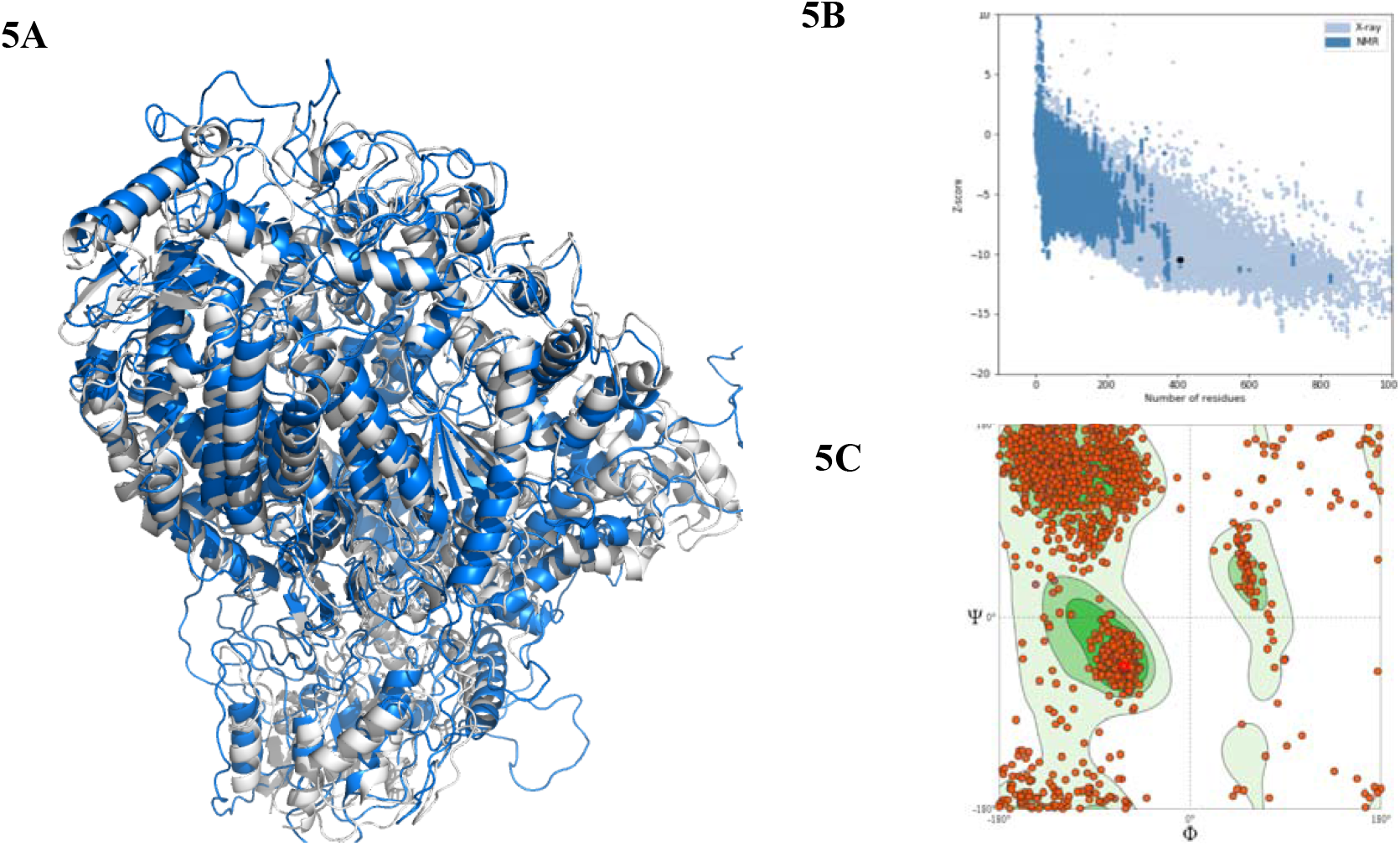
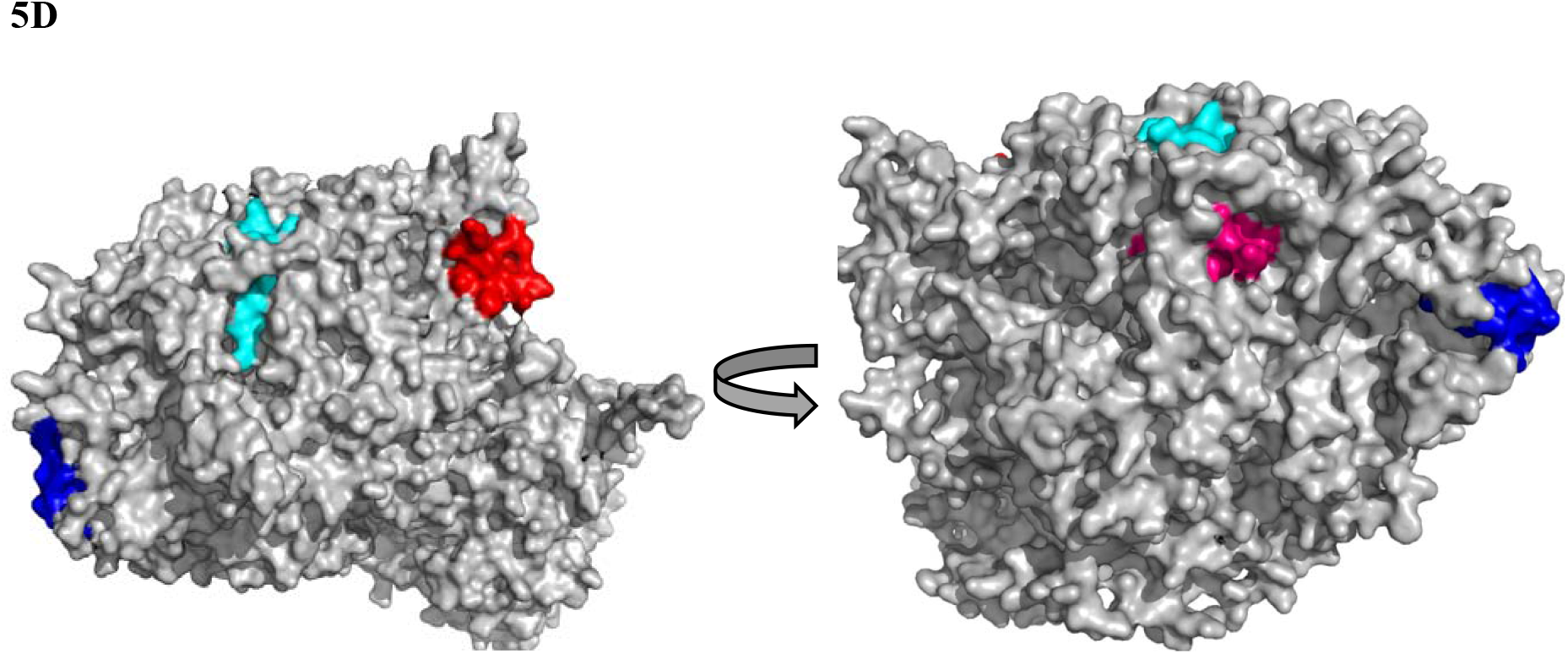
The three-dimensional model of L protein with structural validation and the superficial localities of the predicted epitopes. (**5a**) The predicted model superposed view (grey color represents the model and the blue one is the template). (**5b**) The ProSA Z score for model quality estimation and (**5c**) the Ramachandran plot of the predicted model. In figure **5d** the proposed epitopes SRIKSIKNLSDLTFL (red), KYNLAFRYEFTRHFI (cyan), HHIYRLRNSTALKTF (blue), and LPYRVRNVQTLAEAL (pink), having 9-mer core epitope NLSDLTFLI, FRYEFTRHF, YRLRNSTAL, and YRVRNVQTL, respectively are shown.

### 3.5 Molecular docking and association of HLA allele’s scrutiny

The structural minimization of the peptide was done before the docking analysis and the minimization obtained from energy level 62.3 to −523.53 for 15-mer and −287 to −539.43 for 9- mer, respectively. The 9.0-mer (YRLRNSTAL) and its 15-mer (HHIYRLRNSTALKTF) form of the epitope were docked in the furrow of the HLA-C*07:02 and DRB1*04:01 with docking score of −8.6 and −6.3 kcal/mol, correspondingly. Auto Dock Vina developed various configurations of the docked peptide and picked the better one at an RMSD (root-mean-square deviation) score of 0.0 for the ultimate measurement. The PyMOL (Version 1.5.0.4) was used for the visualization of the docking poses. The amino acid residues- Glu-63, Lys-66, Gln-70, Ser-77, Asp-114, Thr-143, and Gln-155 form hydrogen bond (**Figure 6)** with the 9.0-mer epitope and the residues- Tyr-30(B), Asn-62(A), Asp-66(B), Gln-70(B), Lys-71(B) and Thr-77(B) form hydrogen bond (**Figure 7)** with the 15.0-mer. In addition, the control docking energy for MHC-I interaction was found to be −9.6 kcal / mol and is shown in **Figure S2**.

**Figure 6:**
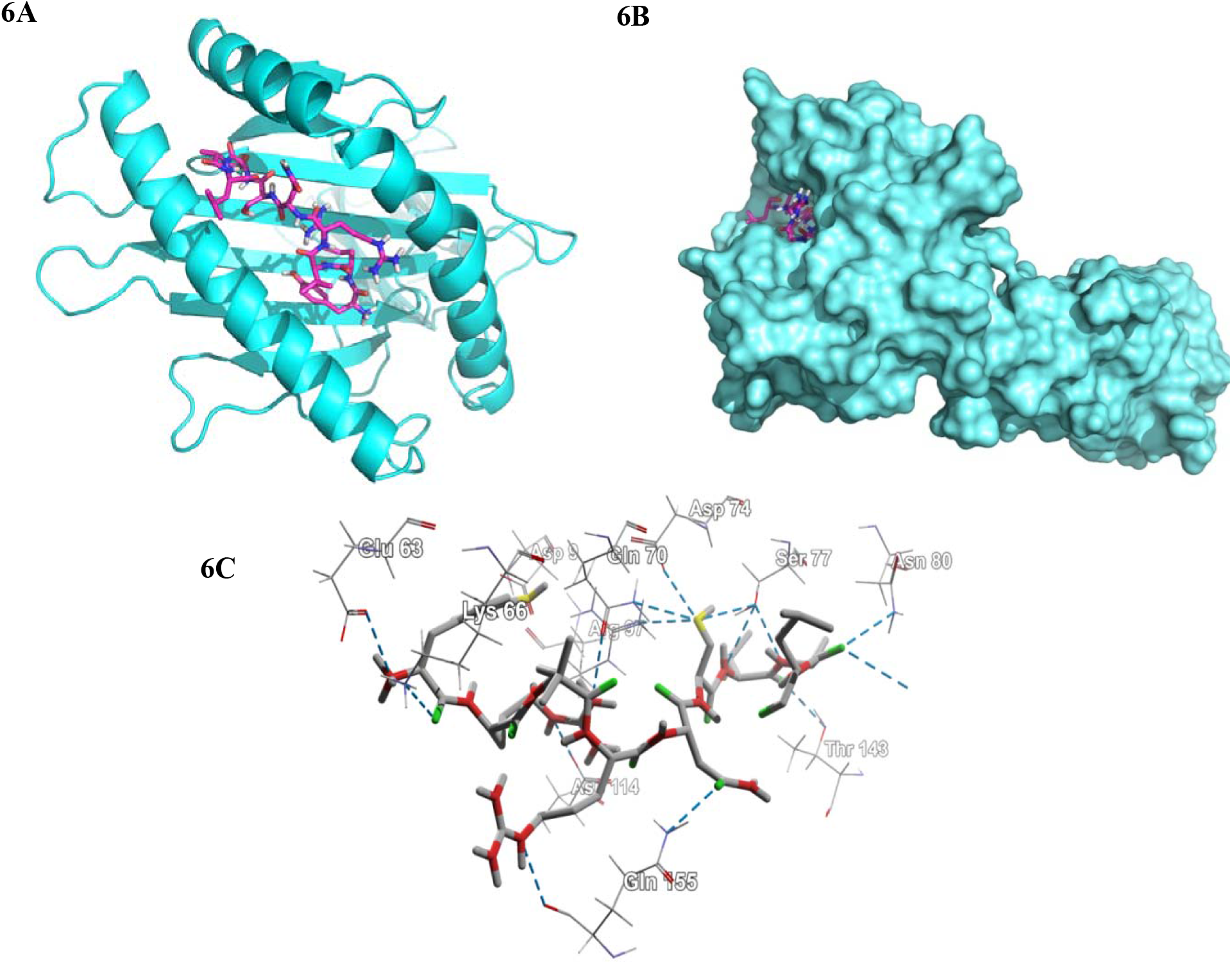
Docking analysis of the predicted epitope YRLRNSTAL and HLA-C*07:02. (**6a**) Representing the cartoon view. (**6b**) Representing the surface view of the interaction and convincing the perfect binding. (**6c**) Representing the interacted amino-acid residues with the peptide.

**Figure 7:**
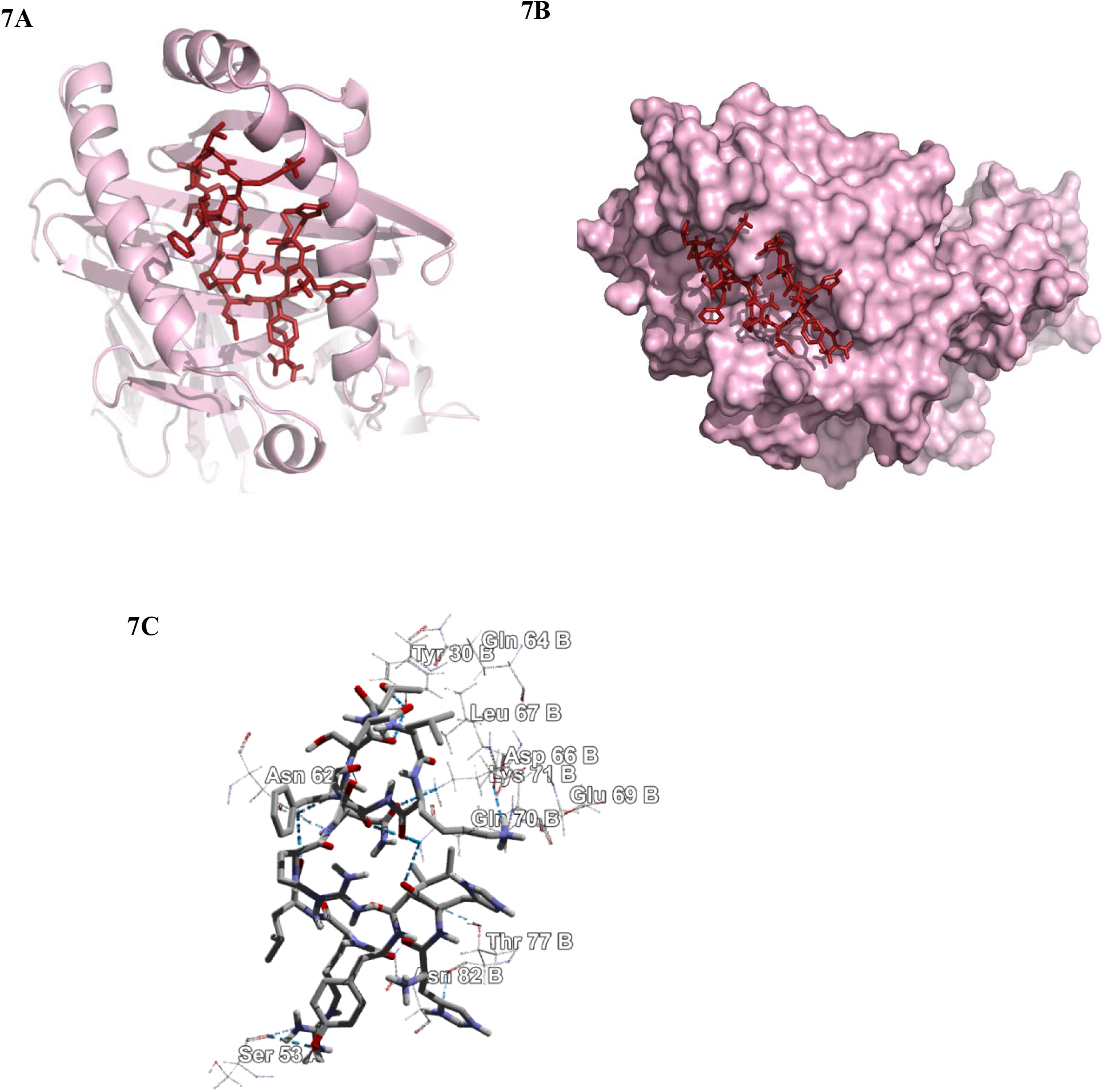
Docking analysis of the predicted epitope HHIYRLRNSTALKTF and HLA- DRB1*04:01 allele. (**7a**) Representing the cartoon view. (**7b**) Representing the surface view of the interaction and convincing the perfect binding. (**7c**) Representing the interacted amino-acid residues with the peptide.

### 3.6 Allergenicity Assessment and B-Cell Epitope Detection

AlgPred predicted allergenicity of the epitope depending on amino acid structure. The predictive score of AlgPred for the two combined epitopes was −0.51324781, with a cutoff value −0.4.

The linear peptide with 15-mer form (HHIYRLRNSTALKTF) through the sequence-based methodology, the B-cell epitope calculation was accomplished and the different prediction parameters were considered and the values ranging from −0.929 to 3.214. The cutoff values for these predictions were ranging from 0.500 to 1.390 (**Figure 8**). The maximum antigenicity score of the peptide calculating through Kolaskar and Tongaonkar antigenicity scale was 1.074 (**Figure 8a**). Accessibility of peptide surfaces is another crucial benchmark for satisfying the criteria of a prospective B-cell epitope and here the Emini surface accessibility calculation supports the prediction with a maximum value of 1.871 (**Figure 8d**). The Parker hydrophilicity prediction was also employed with a maximum score of 3.214 and is shown in **Figure 8f.** Additionally, the maximum score of the Chou and Fasman beta turn estimation score was 1.083 (**Figure 8c**), the flexibility estimation score of Karplus and Schulz was 1.066 (**Figure 8e**), and the Bepipred linear epitope prediction analysis was 0.601 (**Figure 8b**).

**Figure 8:**
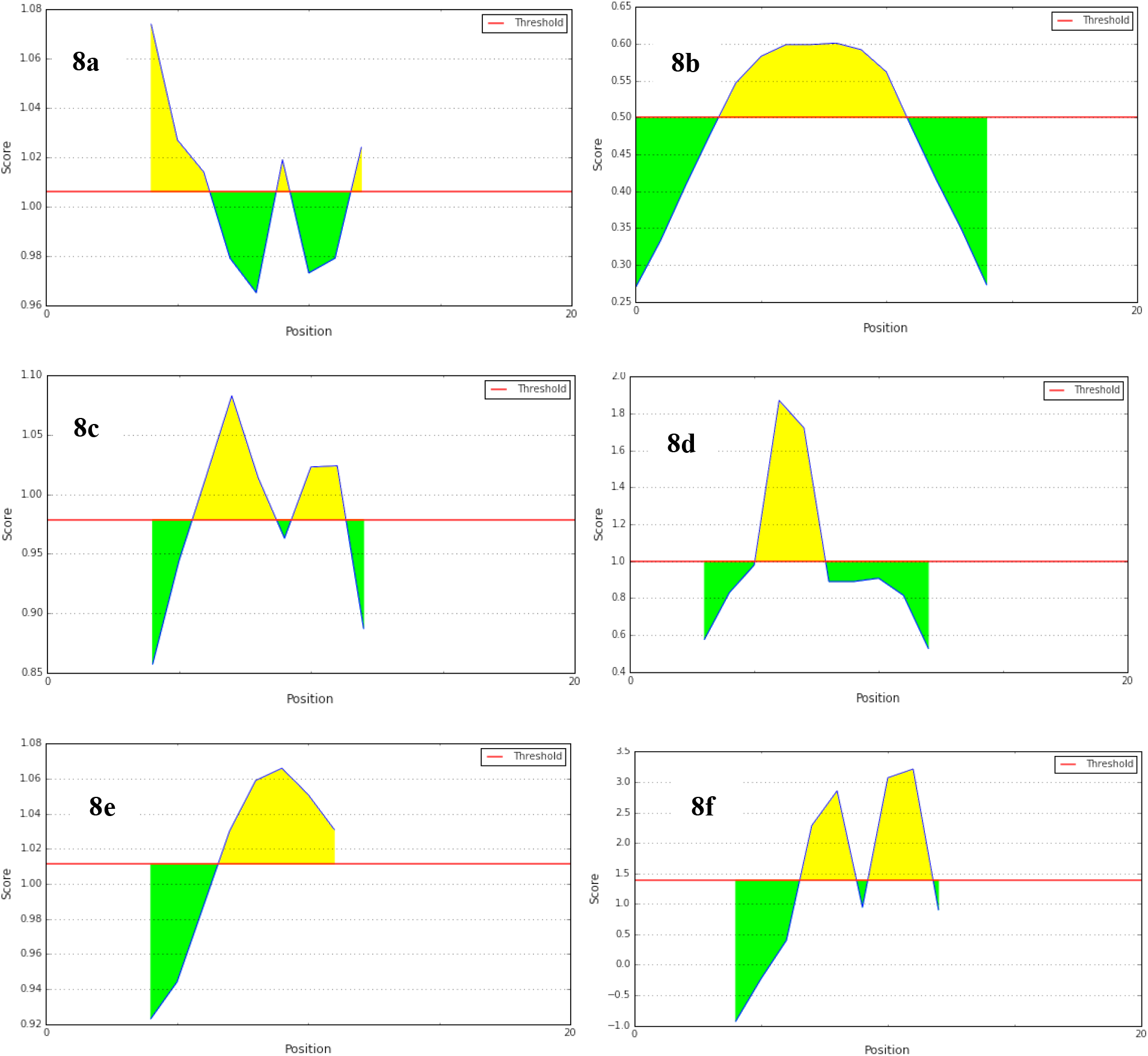
B-cell epitope prediction. (a) Kolaskar and Tongaonkar antigenicity prediction of the proposed epitope with a threshold value of 1.06. (b) Bepipred linear epitope prediction of the proposed epitope, with a threshold value of 0.50. (c) Chou and Fasman beta-turn prediction of the epitope, with a threshold of 0.98. (d) Emini surface accessibility prediction of the epitope, with a threshold of 1.1. (e) Karplus and Schulz flexibility prediction of the epitope, with a threshold of 1.1. (f) Parker hydrophilicity prediction of the epitope, with a threshold of 3.2. Notes: the x-axis and y-axis represent the sequence position and antigenic propensity; respectively. The regions above the threshold are antigenic (desired), shown in yellow.

## 4.0 Discussion

Although the epitope-based design of vaccines has become a conjoint method, no substantial research has yet been done in the case of the Marburg virus L-protein. Marburg virus genome is composed of ribonucleic acid rather than deoxyribonucleic acid. It is especially challenging to develop vaccines for RNA viruses due to rapid mutations of various surface proteins [49]. Therefore, targeting transcription or replication machinery is the possible way to establish effective antiviral therapies against RNA viruses like the Marburg virus. Scientists have discovered that L-protein is a significant cellular element for Marburg virus genome transcription and duplication. Once a cell is infected by the Marburg virus, its genetic RNA code goes into the cell accompanied by L-protein. The L-protein usually “reads” the RNA code and produces messenger RNA that creates viral proteins along with duplication and the development of additional viral constituents. For the above mentioned significant contributions, the L-protein was selected to scheme the utmost plausible epitopes by computational methodologies.

Sequence-based bioinformatics methods have been used to model epitopes of both B cells and T cells to impart immunity in distinct manners. Many vaccines are currently constructed on immunity from B cells; but recently vaccinations depending on the T-cell are widely adopted. Antigenic drift can quickly circumvent antibody mediated reactions over durations, whereas cellular immunity also delivers long-term protection [50, 51]. The cytotoxic CD8+ T lymphocytes (CTL) prevent the dissemination of contagious agents through detecting and killing affected cells or releasing exclusive cytokines specific for virus. [16, 52]. Thus, vaccination based on T-cell is a distinctive method for obtaining a robust protective response counter to viruses [14, 53].

From the phylogenetic analysis, it was quite clear that all the different L-proteins have different origins and having diverse groups in the phylogenetic tree (**Figure 2**). We have also identified the divergence among the sequences (**Figure S1**). From that point of view, we selected the most antigenic protein, AFV31373.1, which suggested its ability to elicit a potent immune response.

NetCTL server was used to find 48 epitopes from 12 MHC supertypes (**Table 1**), and initially, we selected 26 to activate the T-cell response. The epitopes NLSDLTFLI, FRYEFTRHF, YRLRNSTAL, and YRVRNVQTL are principally chosen to scheme a vaccine based on the primary analysis, including the attraction with MHC class I and combined score processed by NetCTL server (**Table 1** and **2**). The conservancy is an epitope’s one of the most critical criteria for selecting it for the production of vaccines [14]. For our proposed epitopes, conservancy study showed conservatives of 87.23, 100, 94.74, and 100 percent, respectively, among all available sequences (**Table 4**). The locations of the four epitopes anticipated are shown in MSA of L proteins in **Figure 3**.

Vaccine candidates should have superior population coverage to achieve acceptability [14]. Before designing, that is very important. In our analysis, we found that the combined population coverage of our proposed epitopes was 93.48, 93.18, 94.42, and 91.33 percent, respectively. This outcome revealed that the proposed epitopes have broader in vitro coverage; therefore, they would be supreme candidates for consideration of vaccines.

For visualizing the precise position of the anticipated epitopes, the tertiary structure of the targeted protein was generated and authenticated by Ramachandran Plot (**Figure 5**), whereby 90.51% of amino acid molecules were found within the preferred zone. The Z scores from the ProSA server for the protein were −8.96, which also support the validity of the predicted models, as the values were within the plot and close to zero. That supports reasonably good model, as there is currently no available crystal structure for the L-protein of MABV. As the epitopes were present on the model’s surface (**Figure 5**), the chance of interaction with the immune system will be improved as early as possible [14, 15].

Lastly, epitopes YRLRNSTAL and YRVRNVQTL (15.0-mer length, HHIYRLRNSTALKTF and LPYRVRNVQTLAEAL, respectively) were identified as being the utmost potent and greatly interrelated HLA aspirants for class II MHC molecules (**Table 3**). Our suggested epitopes are also of a non-allergenic type as per the FAO / WHO Allergenicity Evaluation System, which is another vital peptide vaccine criterion [14, 16, 54].

On the basis of the combined score (2.546), the core epitope YRLRNSTAL would be the paramount epitope applicant, thus further exposed to binding aptitude study. The docking study ensures accuracy with a fairly high binding score and the correctly directed interfaces between the MHC and the predicted epitopes. In addition, comparative research with the empirically observed peptide-MHC complex has showed the specificity of our estimation with comparable binding energy and interacted residues (Figure 6, 7 and **S2**).

Throughout our research, the B-cell epitope and the T-cell epitope were both given preference, which can activate both primary and secondary antibody mediated immune response [13, 55]. Multiple methods of prediction were used to decide the B-cell epitope, taking into account numerous benchmarks of antigenicity, beta-turn, hydrophilicity, surface accessibility, and two other methods. Our proposed epitope complied with all the criteria of the B-cell prediction techniques mentioned earlier (**Figure 8**).

In conclusion, we are clearly optimistic from all of the above *in silico* studies that our recommended epitope will cause an immune response in vitro and in vivo.

## 5.0 Conclusion

Prevention and monitoring during outbreaks of recently-evolving Marburg virus contagions are both necessary and challenging. Through *in silico* analyses, the laboratory experimental work can be guided by finding the desired solutions with less tests and recurrences of errors, hence saving the researchers’ time as well as costs. We used different computational methods in this analysis to identify a possible epitope against the Marburg virus. Our vaccinomics approaches speculate that the selected part of the L-protein is a promising applicant for a peptide vaccine. However, to validate the effectiveness of the deduced amino acid sequences as an epitope vaccine in case of the deadly Marburg virus, further wet laboratory justification is required.

## 6.0 Acknowledgements

None declared.

## 7.0 Author contributions

Tahmina Pervin conceived, designed, carried out the analysis, and drafted the manuscript. Arafat Rahman Oany has directed the report, assisted in the writing, review and crucial revision of the manuscript. The final Manuscript was read and accepted by both contributors.

## 8.0 Confliction declaration

None.

## 9.0 Ethics approval

Not Applicable

